# Forward genetics by sequencing EMS variation induced inbred lines

**DOI:** 10.1101/045427

**Authors:** Charles Addo-Quaye, Elizabeth Buescher, Norman Best, Vijay Chaikam, Ivan Baxter, Brian P. Dilkes

## Abstract

In order to leverage novel sequencing techniques for cloning genes in eukaryotic organisms with complex genomes, the false positive rate of variant discovery must be controlled for by experimental design and informatics. We sequenced five lines from three pedigrees of EMS mutagenized Sorghum bicolor, including a pedigree segregating a recessive dwarf mutant. Comparing the sequences of the lines, we were able to identify and eliminate error prone positions. One genomic region contained EMS mutant alleles in dwarfs that were homozygous reference sequence in wild-type siblings and heterozygous in segregating families. This region contained a single non-synonymous change that cosegregated with dwarfism in a validation population and caused a premature stop codon in the sorghum ortholog encoding the giberellic acid biosynthetic enzyme ent-kaurene oxidase. Application of exogenous giberillic acid rescued the mutant phenotype. Our method for mapping did not require outcrossing and introduced no segregation variance. This enables work when line crossing is complicated by life history, permitting gene discovery outside of genetic models.This inverts the historical approach of first using recombination to define a locus and then sequencing genes. Our formally identical approach first sequences all the genes and then seeks co-segregation with the trait. Mutagenized lines lacking obvious phenotypic alterations are available for an extention of this approach: mapping with a known marker set in a line that is phenotypically identical to starting material for EMS mutant generation.

## INTRODUCTION

One of the most effective and convincing methods to demonstrate gene function is the molecular identification of allelic variation responsible for induced mutant phenotypes. This gene function discovery approach, originally known as genetics and now sometimes referred to as “forward genetics”, has dramatically expanded our understanding of development, physiology, and biochemistry over the last century (Rice, 2014). Much of this understanding has been generated in genetic model systems, which are characterized by simple laboratory culture, short generation times and small genome sizes, but limited in their physiology, anatomy, and biochemistry (Hunter, 2008; Trontin et al., 2011). Historically, identification of the genes responsible for mutant phenotypes was accomplished by map-based cloning. Traditional map-based cloning relies on outcrossing, which introduces phenotypic diversity that can confound mutant scoring. Mutant phenotypes with dramatic differences and insensitive to variation introduced by the crossing partner are the most amenable to map-based cloning approaches. Linkage of the mutant phenotype to a chromosomal region is used to generate a list of candidate causative genes. Recombinant genotypes sufficient to recapitulate the mutant phenotype are then used to narrow the candidate gene list. Mapping genetic loci with sufficient precision to identify a single locus requires ruling out polymorphisms introduced by the outcrossing partner and sequencing a limited number of candidates to identify a single remaining causative polymorphism.

While the number of polymorphisms in a mutagenized genome may scale with the overall genome size, the number of genome positions capable of affecting a change in a phenotype does not. EMS (ethyl methanesulfonate) mutagenesis induces a few thousand polymorphisms per individual; a relatively small number when compared to the hundreds of thousands that differ between diverse lines. Unlike insertion mutagenesis (e.g. transposons or T-DNA) or natural variation, chemical mutagens induce changes in phenotype by modifying protein-coding capacity and only very rarely are non-coding sequence mutations in cis-regulatory sequences identified as causative mutations. Essentially, a genome’s coding capacity determines the phenotypically-affective mutational target. For example, in a genome with 30 Mbp of protein coding sequence, a 1/125,000 bp mutation rate would correspond to ~240 genic mutations. As approximately one third of all EMS alleles result in synonymous changes (Ashelford et al., 2011; Henry et al., 2014; Thompson et al., 2013) this would mean 160 potentially causative non-synonymous mutations per line. EMS is an effective way to induce mutations in protein-coding sequence, which results in an observable phenotype.

Most next generation sequencing (NGS) applications utilize large segregating populations to clone genes by bulked segregant analysis (Michelmore et al., 1991). This has been demonstrated in a variety of species including Arabidopsis, Tef, rice, zebrafish, sorghum, and drosophila (Abe et al., 2012; Austin et al., 2011; Lindner et al., 2012; Mokry et al., 2011; Schneeberger et al., 2009; Uchida et al., 2011; Zhu et al., 2012a; Zhu et al., 2012b; Rizal et al., 2015). In organisms in which extensive pedigree data is often kept, as is the case for many crops and model systems, subtraction of pre-existing variation, and variants present in unaffected siblings would allow the rapid elimination of large genome sections as possible sites for mutant loci. In self-fertile organisms, recurrently selecting heterozygous individuals and selfing them to make advanced single lineage families can drive all loci but the mutant locus to homozygosity. This creates affected and unaffected siblings that vary for the single causative polymorphism only. Such an approach leverages the polymorphism discovery by NGS and the use of pedigree information to identify the causative polymorphism.

Sorghum is a self-fertile diploid panicoid grass that includes crops for food, forage, and biomass products. The adaptation of sorghum to hot and arid environments makes improved sorghum a major contributor to global food security. The sorghum genome of accession BTx623 was sequenced (Paterson et al., 2009). Substantial genetic variation in sorghum accessions has been described (Casa et al., 2008; Hamblin et al., 2004; Mace et al., 2013; Nelson et al., 2011). A number of induced variation collections have also been generated (Blomstedt et al., 2012; Xin et al., 2008). The fully sequenced genome, diploid inheritance, moderate genome size, and availability of induced mutants makes sorghum an excellent system to demonstrate mutant gene identification in complex eukaryotes and organisms of human importance. This would expand the reach of these advanced applications of sequencing and mutagenesis technologies beyond genetic model systems.

Next-generation, short read, sequencing technologies are error prone and this can produce a number of artifacts. These errors, when summed over the length of a complex eukaryotic genome, can result in incorrect inference. Controlling the sources of these errors in the analysis of whole genome sequencing results may lead to better gene mapping and cloning procedures. Some of these errors result from the inherent error rate in the sequencing technology (Minoche et al., 2011; Quail et al., 2008; Quail et al., 2012) and these problems can be overcome by sequencing genomes at greater depth. Other artifacts differentially affect specific positions making them error prone, including alignment errors at insertion or deletion alleles (R. Li et al., 2009), SNPs resulting from mistakes in the reference assembly, variation in research material relative to the reference assembly, poor alignment performance (Cheng, Teo, & Ong, 2014; Fredman et al., 2004; O’Rawe et al., 2013), alignment to repetitive or paralogous sequences (Cheung et al., 2003; Estivill et al., 2002), and the alignment of sequence from DNA encoded by unassembled genomic regions to an incomplete digital reference genome (Teo, Pawitan, Ku, Chia, & Salim, 2012). When mutagenesis only induces a few thousand true changes to the genome, the number of false positives can be far greater than the number of mutations. Methods to remove these errors are needed to make gene identification from mutants of complex eukaryotes to become routine and reliable.

In this work, we compared the results of whole genome, short-read sequencing of a sorghum dwarf mutant to similar data from independently-derived EMS-mutagenized lines. By collecting whole genome sequence data from two phenotypically unaffected individuals in an EMS population and using the data from a previously published unrelated mutant in *dhurrinase2* (Krothapalli et al., 2013), we were able to identify nucleotide positions in the sorghum genome that resulted in SNP calls in more than one lineage. These positions cannot be the causative mutation for any phenotypes unique to one lineage. Initial SNP calls were not consistent with the result of guanine residue alkylation, the mechanism of EMS mutagenesis. After removal of the shared alleles identified between the three mutants and considering only the coding sequence SNPs, the proportion of G:C to A:T changes, the expected changes from EMS treatment, ranged from 79 to 91 percent per line. We measured allele frequencies of these SNPs in sequence data from wild type, heterozygous and, dwarf siblings. The vast majority of these SNPs displayed allele frequencies incongruent with the mutant allele status of the line. Allele frequencies for one region on chromosome 10 segregated with the mutant phenotype, identifying this region as encoding the dwarf mutant. One, and only one, coding sequence change was present in the delimited interval in the dwarf, missing from the wild-type sibling, and heterozygous in the heterozygote sibling’s genome. This polymorphism introduces a premature stop codon in the ent-kaurene oxidase gene of sorghum (Sobic.010G172700), encoding a putative enzyme in gibberellin biosynthesis. The mutation co-segregated with the dwarf phenotype, which was reversed by GA application, and confirmed the causative nature of this polymorphism. Using EMS-induced SNP variation as molecular markers permits any phenotypically unaffected line to act as an EMS Variation Induced Lines (EVIL twins) for mapping crosses without introducing the segregation variation of a distinct and diverse genetic background. The use of pre-existing data to remove error prone positions to improve SNP calling and EVIL twins for mapping should improve both the accuracy and sensitivity of forward genetics.

## RESULTS

### Observation of a dwarf mutant segregating in an M4 generation

Sorghum BTx623 seeds were mutagenized in a prior study and the resulting offspring selfed for three generations to generate M4 individuals (Xin et al., 2008). We screened a limited number of families for aphenotypic EMS-treated derivatives. Two M4 individuals from independent lineages, lines 10 and 12, were selected because they had no obvious mutant phenotypes. When these lines were selfed, the progeny of line 12 were all normal while the M5 progeny of line 10 segregated for extreme dwarfism with reduced leaf lengths and compact internodes (Figure 1). Multiple tall M5 siblings were self-pollinated and M6 families were grown and scored for the dwarf phenotype. Dwarfism segregated as a monogenic recessive trait. One M6 family, the selfed progeny of M5 individual 10-2, was replanted and determined to be a homozygous wildtype derivative of line 10. Another M6 family, the selfed progeny of M5 individual 10-3, segregated 3:1 for tall:dwarf and indicated that individual 10-3 was a heterozygote. Figure 1 shows the pedigree and dwarf phenotype expression of the sequenced lines. We used these materials to test our approach to clone EMS-induced mutants by using all of the EMS-induced variants as molecular markers and build the bioinformatic analysis pipeline necessary to link NGS and genetic mapping.

**Figure 1.**
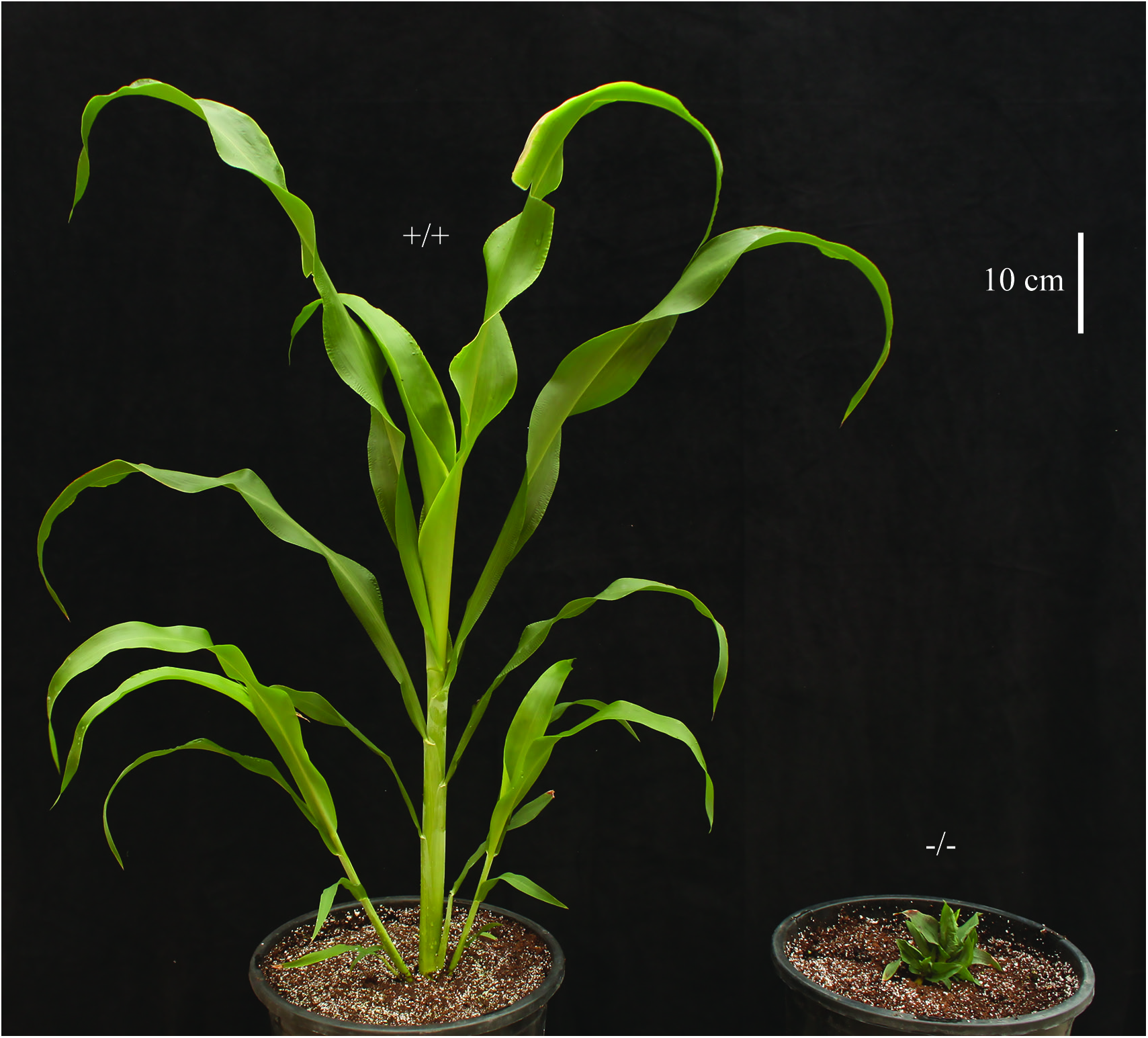
(a). BTx623 (wild type) and 10-d mutant plants, exhibiting a dwarf phenotype. (b) Pedigree of the sequenced EMS-treated lines. Independent mutagenesis resulted in the *dhr2-1* mutant, the 12-2 line and the 10 line. Selfing of 10 resulted in sublines 10-2, 10-3, and 10-d.

### Sequence data generation and read mapping

We sequenced individuals derived from line 10 based on their segregation for the extreme dwarf phenotype. DNA was isolated from a pool of dwarf mutant tissue designated 10-d, a pooled sample of tall heterozygous siblings (the M5 10-3 individual), and a poll of homozygous wild-type sibling from line 10-2. The DNA from each sample was sequenced. In addition, we sequenced another line, 12-2, which showed no gross morphological defects and was independently derived from the same mutagenesis experiment as line 10 (Xin et al., 2008). Finally, we re-analyzed the sequencing data used to clone the *dhurrinase2* (*dhr2-1*) EMS-induced sorghum mutant, which was found in a different mutant population (Krothapalli et al., 2013). A total of 1,036,765,083 Illumina NGS reads were generated from DNA samples derived from the five sorghum EMS-mutant individuals (Table 1). Using the BWA short read aligner (Li and Durbin, 2009), 91% (942,006,177) of the sequenced reads mapped to the reference BTx623 sorghum genome (Paterson et al., 2009). Using the alignment results, the estimated median paired-end insert sizes for the 10-2, 10-3, 10-d and *dhr2-1* sequenced reads were 249, 307, 260 and 297, respectively. The alignment results also showed 88% (889,860,334) of the paired-end reads were properly paired within the range of the above insert sizes. Table 1 shows the summary statistics for the sequencing and alignment to the sorghum genome assembly version 2.1 available at Phytozome (www.phytozome.net). Between 88-94% of the reads mapped to the reference genome. The coverage depths for the homozygous aphenotypic line 10-2, the previously described *dhr2-1*mutant, and the extreme dwarf 10-d were 45-fold, 43-fold and 30-fold genome coverage, respectively (Table 1). Two other individuals were sequenced at lower depths. The aphenotypic 12-2 line and the heterozygous sibling 10-3 individuals were sequenced at 3-fold and 7-fold genome coverage, respectively (Table 1).

**Table 1.**
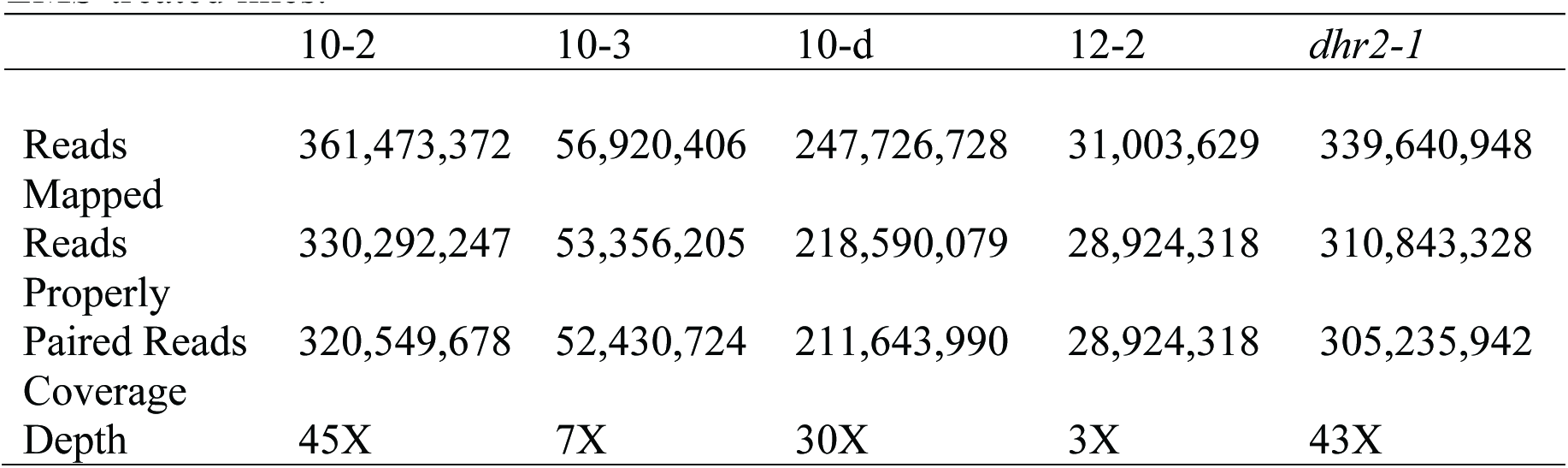
Summary of whole genome sequencing and read mapping for five sorghum EMS-treated lines.

|  | 10-2 | 10-3 | 10-d | 12-2 | *dhr2-1* |
| --- | --- | --- | --- | --- | --- |
| Reads | 361,473,372 | 56,920,406 | 247,726,728 | 31,003,629 | 339,640,948 |
| Mapped Reads | 330,292,247 | 53,356,205 | 218,590,079 | 28,924,318 | 310,843,328 |
| Properly Paired Reads | 320,549,678 | 52,430,724 | 211,643,990 | 28,924,318 | 305,235,942 |
| Coverage Depth | 45X | 7X | 30X | 3X | 43X |

### SNPs detection

We detected SNPs according to a procedure outlined in Figure 2 and described in detail in the methods section. In short, alignments were scored for likely deviation from the reference sequence and then sequentially filtered for coverage, homozygosity, and a requirement that reads map to both strands and sequence data quality. Following this, SNPs were further filtered to remove all variants unlikely to be obtained by EMS treatment, which primarily induces G to A mutations (Greene et al., 2003) which also appear in sequence data as the reverse complement, C to T. Table 2 summarizes the SNP detection results and filtering steps. We obtained a total of 920,834 likely deviations from the reference sequence in initial SNP calling for the five samples using the SAMtools SNP calling package (Li and Durbin, 2009). Discarding low quality SNPs and SNPs of repetitive origin reduced the total by 59 percent to 376,533. Due to the low coverage and depth of sequencing data from the 12-2 sample an unsusually high number of sites initially scored as variants but derived from singleton reads were detected. These positions were efficiently removed by our procedure. Further filtering to retain only SNPs likely to be homozygous with a minimum Phred quality score of 20 retained 125,308 positions, removing 27% of the initial SNP calls. If we further require all reads overlapping a SNP site to contain the mutant allele and for reads to derive from both forward and reverse DNA strands, only 71,058 high-quality homozygous SNPs remain from the five lines. EMS-induced alleles are almost exclusively G:C to A:T transitions unlike background mutations which could be any of the eight possible sequence changes. Removing all but the high quality G:C to A:T SNPs removed 51% of the remaining SNPs. The count of likely EMS-induced changes (high quality G:C to A:T) ranged from 9,668 changes in 10-2 to only 2,635 in the low coverage sequence data from 12-2 (Table 2). An average of 6,914 high-quality G:C to A:T SNPs were identified per sample.

**Figure 2.**
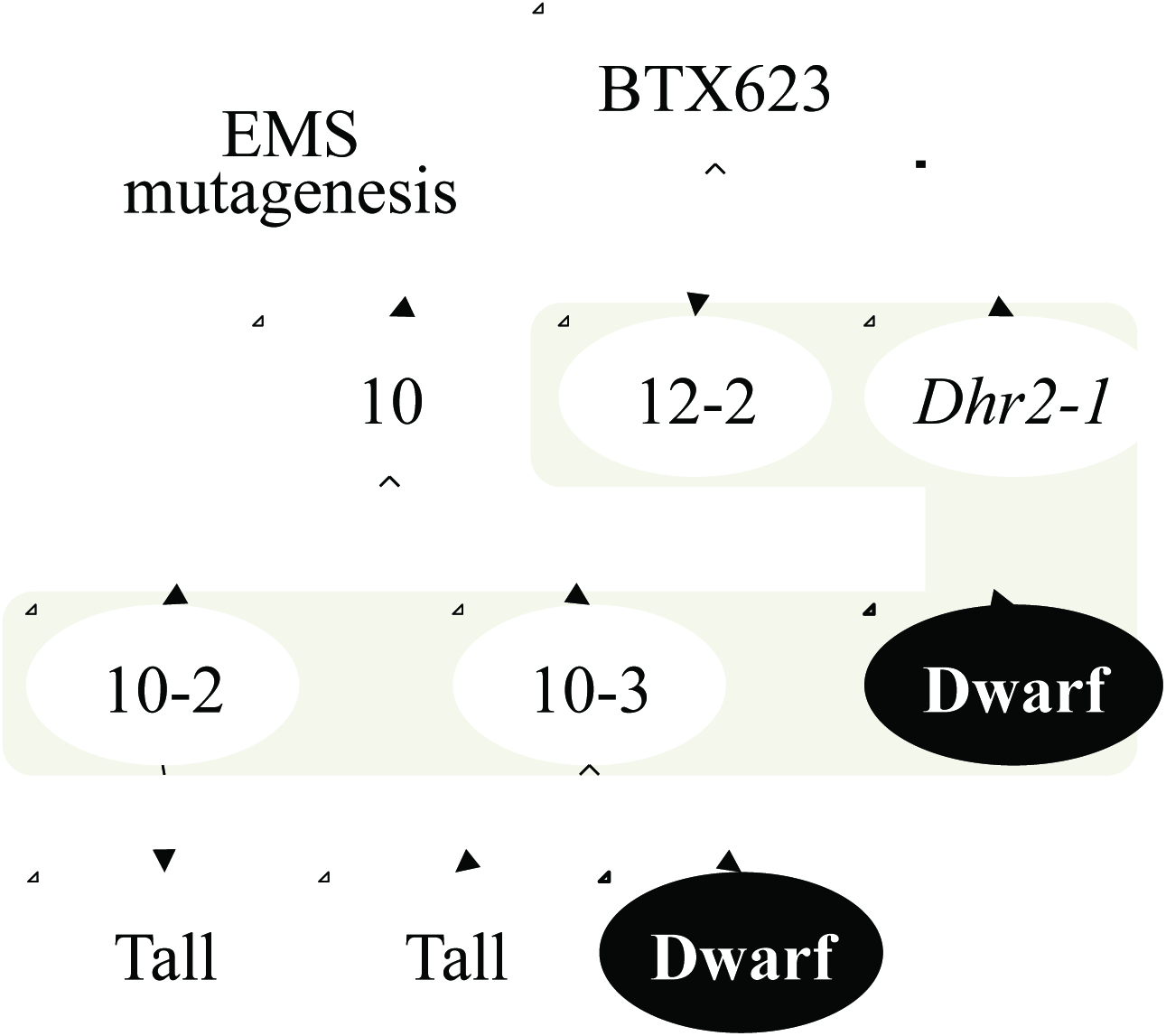
Schematic overview of the SNP detection and annotation pipeline for EMS mutant discovery.

**Table 2.**
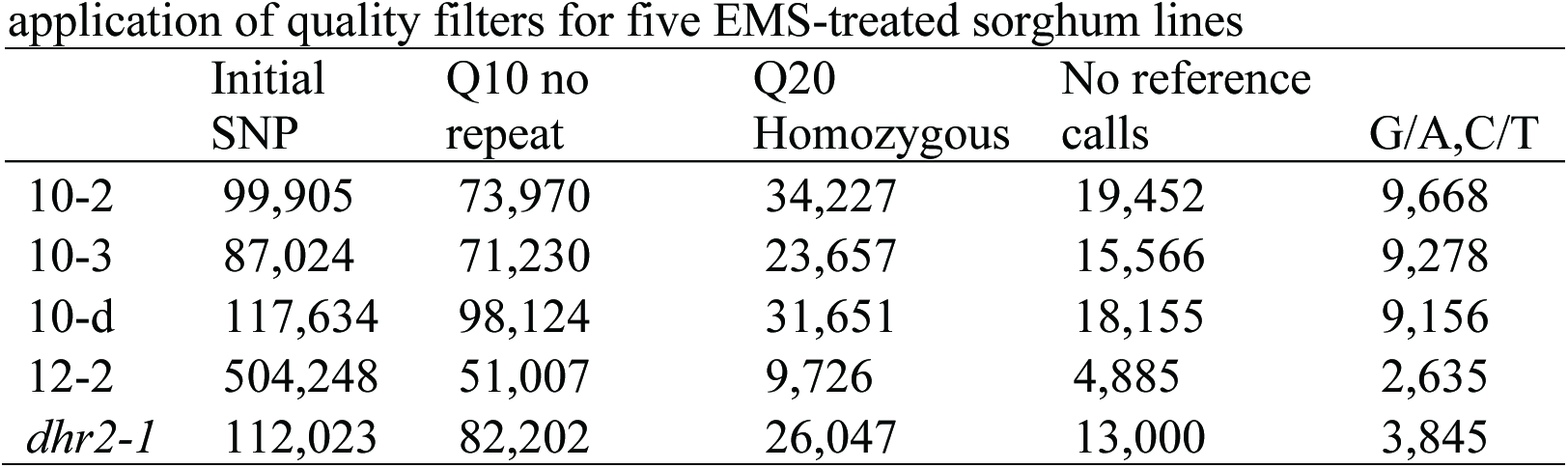
Summary of Single Nucleotide Polymorphism counts during sequential application of quality filters for five EMS-treated sorghum lines

### Independent Mutagenesis as Biological Replication for the Experimental Identification of False Positive SNPs

SNP calling is a statistical likelihood procedure (Nielsen et al., 2011) for each nucleotide position in the genome. As a result, the number of false positives will scale with the overall genome size. Moreover, chemically mutagenized lines have a low true positive rate (1/125,000 to 1/500,000 bp) resulting in a relatively higher proportion of false positives. This is in stark contrast to the use of SNP calling for distinguishing natural populations, in which multiple orders of magnitude more true positives are present. We initially attempted to score each SNP as a molecular marker in our sequencing data and found that many adjacent neighboring SNPs were not linked in the three fully sequenced individuals that shared the “10” pedigree. This violates the predicted inheritance of these chromosomes and DNA segments (data not shown). We hypothesized that a high ratio of false positives in the SNP calling procedure was interfering with mapping and co-segregation. To detect positions in the genome that were false positive SNP calls, we identified genomic positions that were scored as variant in more than one independently mutagenized line. We compared SNP calls from the line 10 derivatives (10-2, 10-3, and 10-d), the aphenotypic EMS-treated line 12 (12-2) and the previously published *dhr2-1* mutant (Krothapalli et al, 2013). SNPs shared between independent lineages are highly unlikely to be due to independently mutagenized chromosomes. Far more likely, any SNP variation shared between the 10, 12, and *dhr2-1* pedigrees would be caused by systematic bias in the generation and analysis of sequencing data. These shared SNPs should detect 1.) errors in the reference sequence, 2.) differences between the mutagenized material and the originally sequenced line, and 3.) positions in the genome that frequently return a likely SNP call due to structure or paralogy. Furthermore, only those SNPs present in the 10 lines and not in 12-2 or *dhr2-1* can be causative for the extreme dwarf locus segregating in that lineage. Figure 3A-C displays Venn diagrams showing the overlap in SNP positions between each of the sequenced line 10 derivatives with the independently mutagenized 12-2 and *dhr2-1* lines. In each case, a large number of SNPs were identified in multiple lineages, indicating that these SNPs are unlikely to be the result of EMS mutagenesis and cannot encode the causative alleles for the extreme dwarf phenotype in line 10. The majority of the shared SNP variation was detected in pairwise comparisons. Only a minority of differences was found when comparing the three lineages of line 10, which indicates the majority of these shared SNPs are not the result of base calling errors in the reference sequence. Table 3 provides the filtered common SNP variations for individual samples considering either genome-wide SNP data or only SNPs in coding sequences.

**Figure 3.**
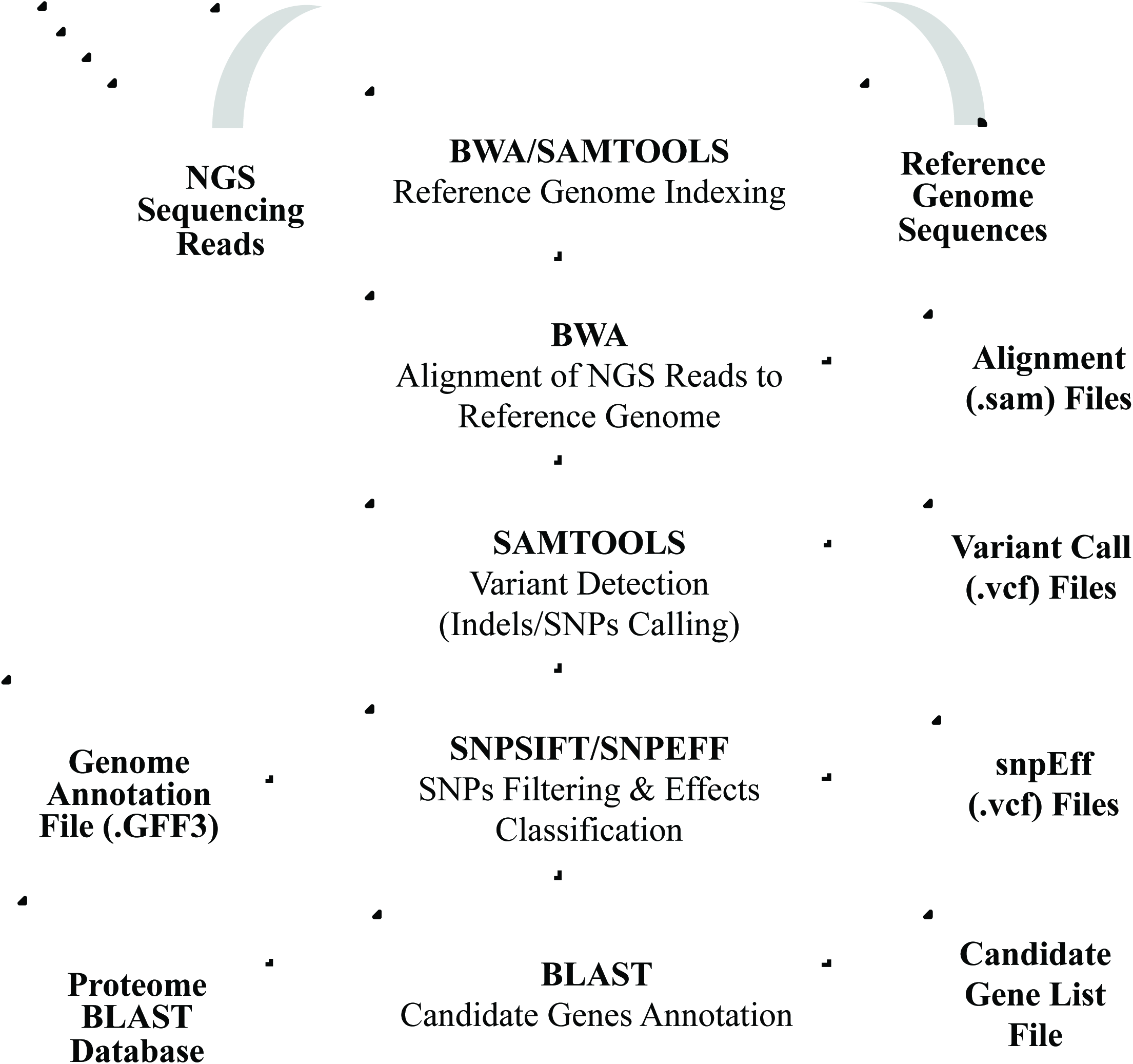
Genome-wide comparison of putative EMS-induced SNPs between pedigrees. Venn diagrams display the number of shared G to A, or C to T SNP positions found in lines 12-2, *dhr2-1,* and (a) 10-2, (b) 10-3, or (c) 10-d.

**Table 3.**
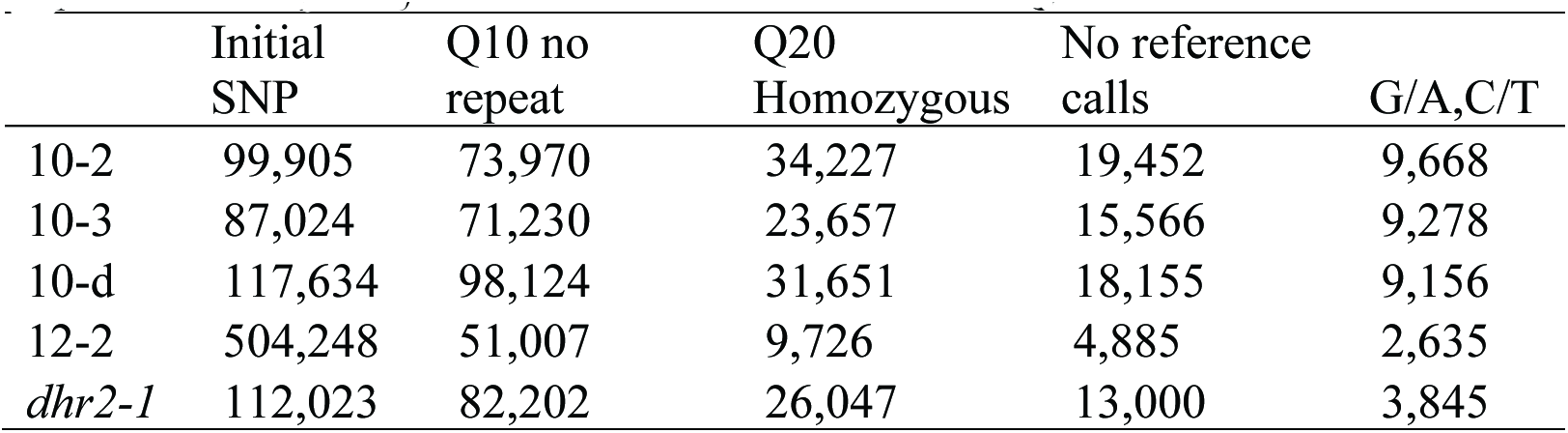
Summary of likely EMS-induced SNP

The subtraction of common variants selectively enriched for EMS-induced G:C to A:T SNPs. In the initial SNP calling, only 20% percent of the SNP calls were G:C to A:T transitions. After quality filtration (previous section; Table 2) 49% of the high-quality SNPs were G:C to A:T variants (Supplementary Table 1). Detection and removal of the SNPs shared between pedigrees resulted in further enrichment with 79% of all lineage-specific SNPs (28,176 SNPs) encoding G:C to A:T transitions (Supplementary Table 1). Thus, roughly one-fifth of SNPs in our stringent sequential SNP filtering procedure were due to false positives arising at reproducible, and therefore detectable, positions in the genome. This proportion was even greater for the coding sequence positions (Table 3) suggesting that paralogy and failed assembly of repeats may contribute to these error prone positions. When considering only the coding sequences, we observed 93% of the SNPs remaining after removal of shared variant positions were G:C to A:T transitions in in the line 10 derivatives (Table 3). This strongly suggests that the overwhelming majority of independently derived high quality SNPs called by this sequential method are homozygous true-positive EMS-induced variants and that the common variants were induced by technical or biological processes distinct from EMS mutagenesis.

### EMS Variant Induced Line mapping of the extreme dwarf mutant to Chromosome 10

If our SNP calling procedure is specific and comprehensive, segregation of the called SNPs within the 10 pedigree should follow Mendelian inheritance and identify the genomic region encoding the dwarf mutant. SNPs linked to the dwarf mutant should be consistently inherited with the recessive trait. SNPs linked to the causative polymorphism will be encoded by their EMS alleles in the 10-d sample, homozygous for the reference allele in the 10-2 wild-type sample, and heterozygous in the 10-3 sample. Figure 4 is a Hormigas plot, so named because each dot representing a SNP resembles an ant, of allele frequencies measured in the wild-type 10-2 at all high-quality G:C to A:T SNP positions identified as homozygotes in 10-d. Only one segment of the 10-2 genome encodes the reference alleles for positions identified as SNPs in the dwarf mutant. This locus on chromosome 10 is predominantly homozygous wild type in 10-2 and located within a region of chromosome 10 conspicuously lacking homozygous 10-d SNP calls, consistent with segregation and recombination in the heterozygous parent that gave rise to the pool of dwarf plants that were sequenced (Figure 4). Thus, based on the sequence data from two libraries, the dwarf mutant can only be located within a relatively narrow window of 5 Mb present on chromosome 10.

**Figure 4.**
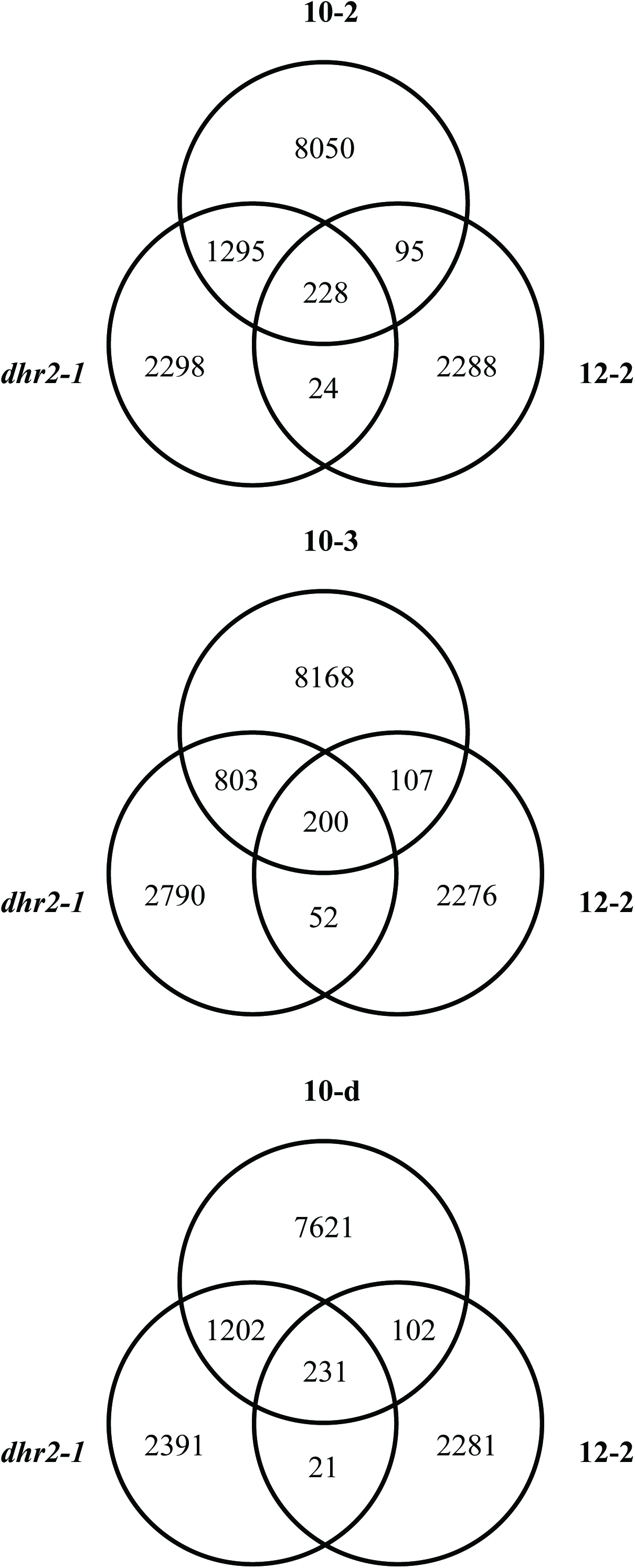
Mapping of the critical region for the 10-d mutant using the EMS mutations in the line and no outcrossing. Hormigas plot of the allele frequencies in the wild-type segregant 10-2 for homozygous polymorphisms detected in 10-d. The only segment not displaying EMS-induced variation is present on chromosome 10.

### The sorghum ent-kaurene oxidase ortholog is responsible for the extreme dwarf phenotype

In a classical map-based cloning procedure, first a mapping interval is identified. Once the effort to identify additional recombinants is greater than the effort of sequencing the genes in the region, coding sequence changes are sought. In our case we already have the sequence of all genes in the genome. Thus, in an effort to identify SNPs likely to alter the line’s phenotype, we determined the protein coding sequences affect by each high-quality SNPs. About 85 percent (23,997) of SNPs were in intergenic regions and 2,113 SNPs were within annotated introns (data not shown). In all five lines, a total of 967 SNPs were predicted to alter the coding capacity of an annotated protein-coding gene. Of these, 927 SNPs were missense mutations, 26 were nonsense (stop codon gain) mutations and an additional 14 encoded stop codon loss or splice site mutations.

A total of 296, 209, and 252 high-quality G:C to A:T SNPs impact protein coding capacity in the 10-2, 10-3 and 10-d sorghum individuals, respectively (Table 3). Of these, only those polymorphisms that are homozygous in the 10-d dwarf mutant and not called in either the wild-type sibling or heterozygous sibling could be the causative polymorphism for the dwarf phenotype. Among the 252 SNPs in 10-d, 154 were shared with both 10-2 and 10-3 and an additional 89 were shared with either 10-2 or 10-3 (Figure 5). This left six SNPs as possible causal EMS-induced mutations for the dwarf phenotype. Of these six, only one sits within the critical window on chromosome 10 identified by allele frequency mapping of the EMS-induced alleles in the wild-type segregant. The detected EMS-induced G to A mutation occurred at genomic location 50,477,994 on chromosome 10 in the sorghum genome (Table 4).

**Figure 5.**
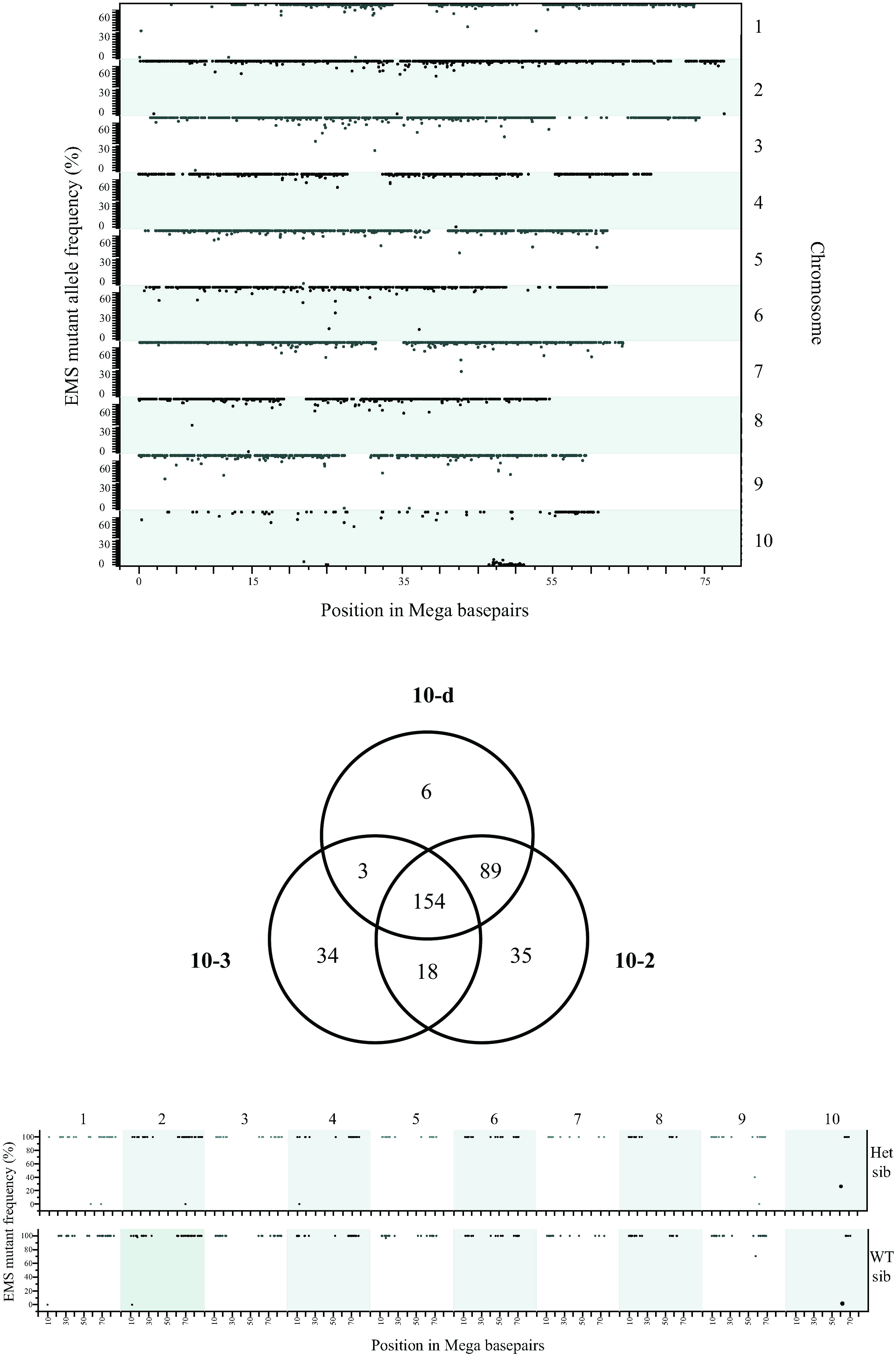
Genome-wide comparison of non-synonymous SNPs present in the lines derived from EMS lineage 10. Venn diagrams display the number of shared G to A, or C to T SNP positions found in lines 10-d, 10-2, and 10-3.

**Table 4.** Putative EMS variants with predicted impact on protein function specific to the dwarf mutant

| Chr | Position | Codon Change | Amino acid change | Gene ID | Gene Description |
| --- | --- | --- | --- | --- | --- |
| 1 | 23070815 | gCc/gTc | A100V | Sobic.001G234400 | Novel |
| 2 | 2006984 | cGg/cAg | R138Q | Sobic.002G021300 | Novel |
| 8 | 52996874 | Gtg/Atg | V13M | Sobic.008G169000 | Novel |
| 9 | 47735642 | Ccg/Tcg | P28S | Sobic.009G124000 | Unknown (DUF3411) |
| 9 | 50971573 | cGg/cAg | R346Q | Sobic.009G153200 | At Plant U-box 29 |
| 10 | 50477994 | Cag/Tag | Q225stop | Sobic.010G172700 | *GA requiring 3* |

We utilized a similar allele frequency analysis focusing on high-quality G:C to A:T SNPs in coding regions to provide additional evidence for the causative polymorphism on chromosome 10. The causative polymorphism for the dwarf mutant will be present as homozygous mutant allele in 10-d, present in intermediate frequency in the reads from the 10-3 heterozygote, and present as the homozygous reference BTx623 allele in 10-2. The allele frequencies from all high quality homozygous G:C to A:T SNPs affecting protein coding changes present in the dwarf mutant, 10-d, are presented as Hormigas plots for both 10-3 and 10-2 in Figure 6. Among all 252 homozygous coding sequence differences identified in 10-d, *exactly one change* was homozygous wild type in 10-3 and heterozygous in 10-2. Again, this corresponded to the G to A mutation at genomic location 50,477,994 on chromosome 10. PCR-based genotyping of this polymorphism was done for twenty-three dwarf individuals derived from 10-2 as well as ninety-seven dwarfs from six independent families from the inital 10 line. There was complete linkage of the dwarf mutant phenotype and the SNP at 10:50,477994 (data not shown).

**Figure 6.**
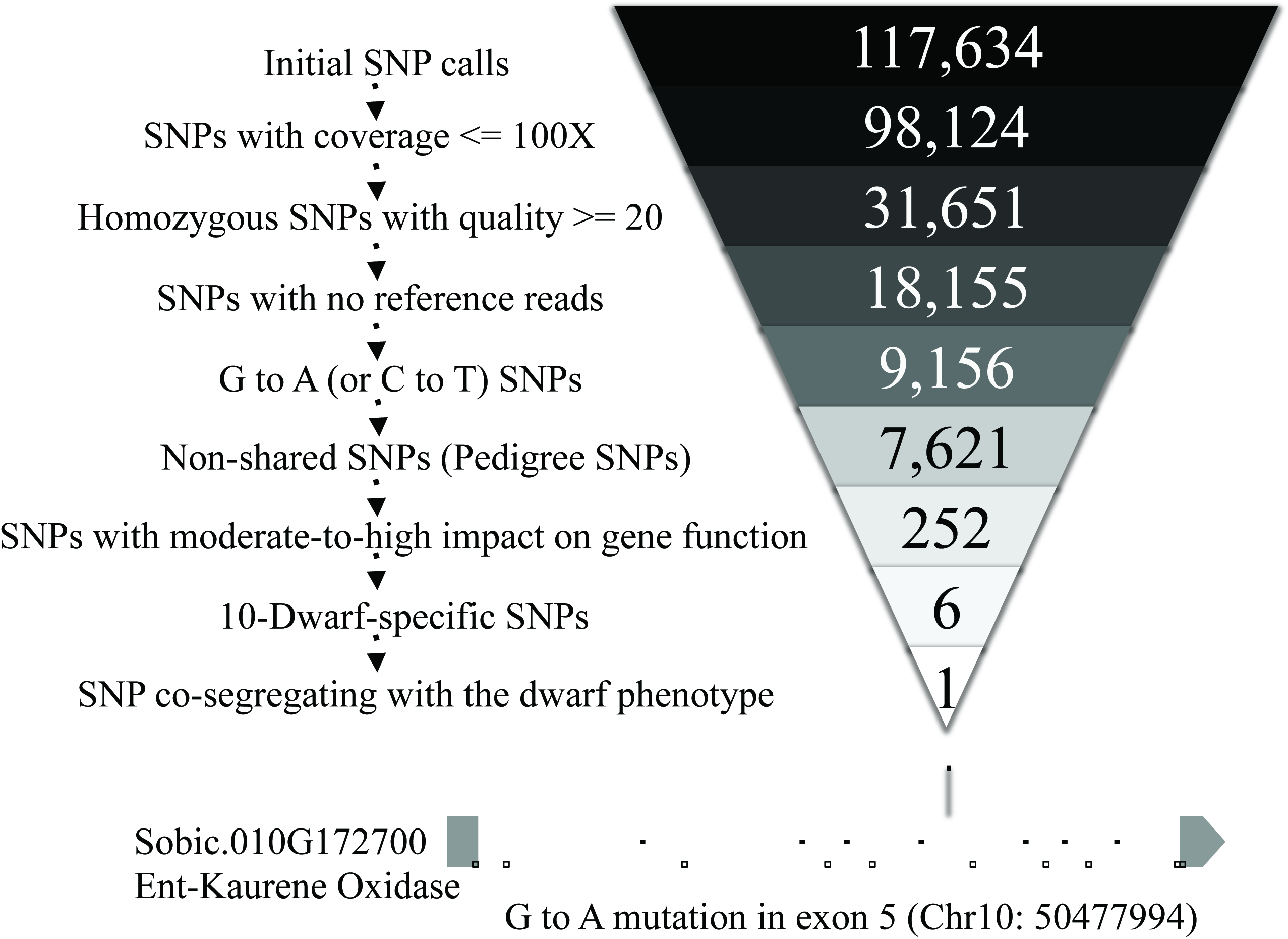
Comparison of allele frequencies for non-synonymous SNPs identifies one and only one SNP that could encode the dwarf mutant. Hormigas plot of the allele frequencies (y-axis) in the heterozygous sibling 10-3 (top panel) and the homozygous wild-type sibling 10-2 (bottom panel) for SNP positions homozygous in the 10-d lineages (x-axis).

The sequential filtering that resulted in the identification of this single SNP is presented in Figure 7. The mutation resides in the sorghum gene Sobic.010G172700 (Table 4), which encodes a 508 amino acid protein, and results in a conversion of the 225^th^ codon from a glutamine (CAG) to an amber stop codon (TAG). Sobic.010G172700 encodes cytochome P450 CYP701A6 and is the sorghum orthorlog of the ent-kaurene oxidases from rice, Arabidopsis, and Pea (Davidson et al., 2004; Helliwell et al., 1999). This enzyme is responsible for multiple reactions in the biosynthesis of the plant hormone gibberlic acid (GA).

**Figure 7.**
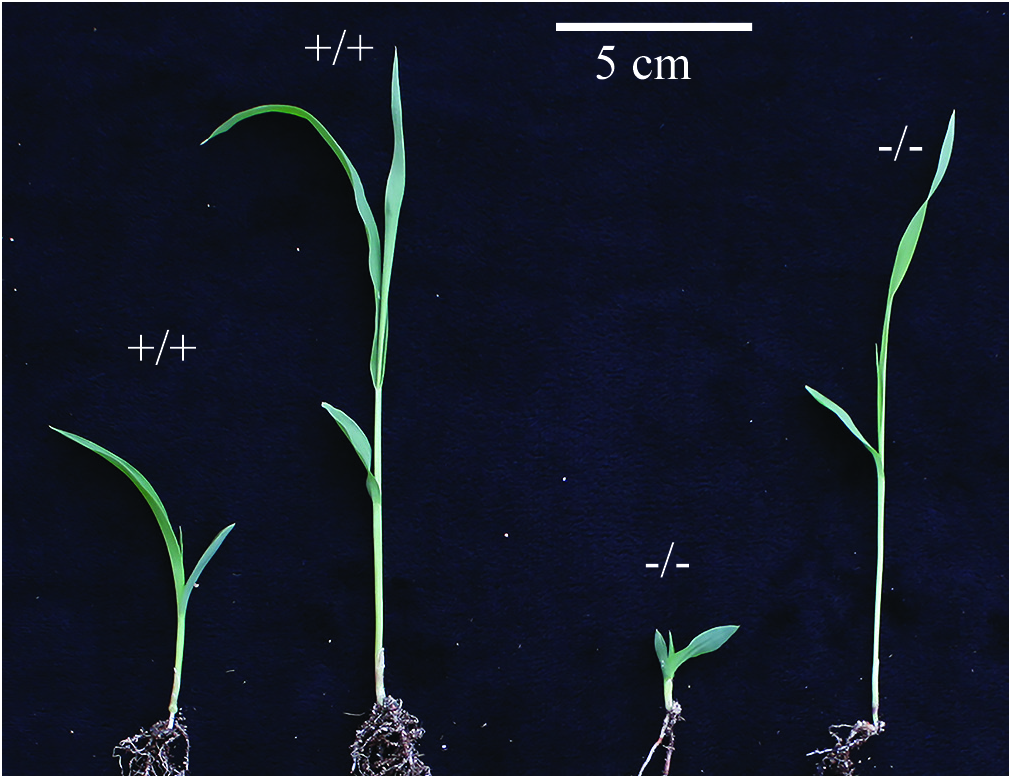
Number of candidate positions after SNP filtering of NGS data for the dwarf mutant in line 10-d. A presentation of the SNP filtering steps are listed on the left and the number of SNPs remaining the genome after each step is provided on the right. The one and only one SNP remaining after filtering is represented graphically. The gene model of the locus responsible for the dwarf phenotype in 10-d is displayed with exons and introns indicated with the position of the mutation marked and annotated.

To confirm that disruption of GA biosynthesis was responsible for the dwarf phenotype, we tested whether the dwarf phenotype could be reversed by the exogenous application of GA3. Application of 10 uM GA3 to light-grown seedlings fully restored wild type growth to the homozygous dwarf phenotype (Figure 8A and 8B). Similarly, the elongation of mesocotyls in dark grown seedlings was restored when dwarf plants were provided 10uM GA_3_ (Figure 8C). Application of GA into to the whorl of light-grown plants substantially rescued the dwarf phenotype in adult plants (data not shown), demostrating that this phenomena extended beyond seedlings. The dwarfism in the 10-d line was reversible with GA_3_ application was consistent with the mutation discovered in Sobic.010G172700 resulting in a disruption in GA biosynthesis. We therefore name the mutation *ent-kaurene oxidase dwarf-1* (*eko-1*).

**Figure 8.**
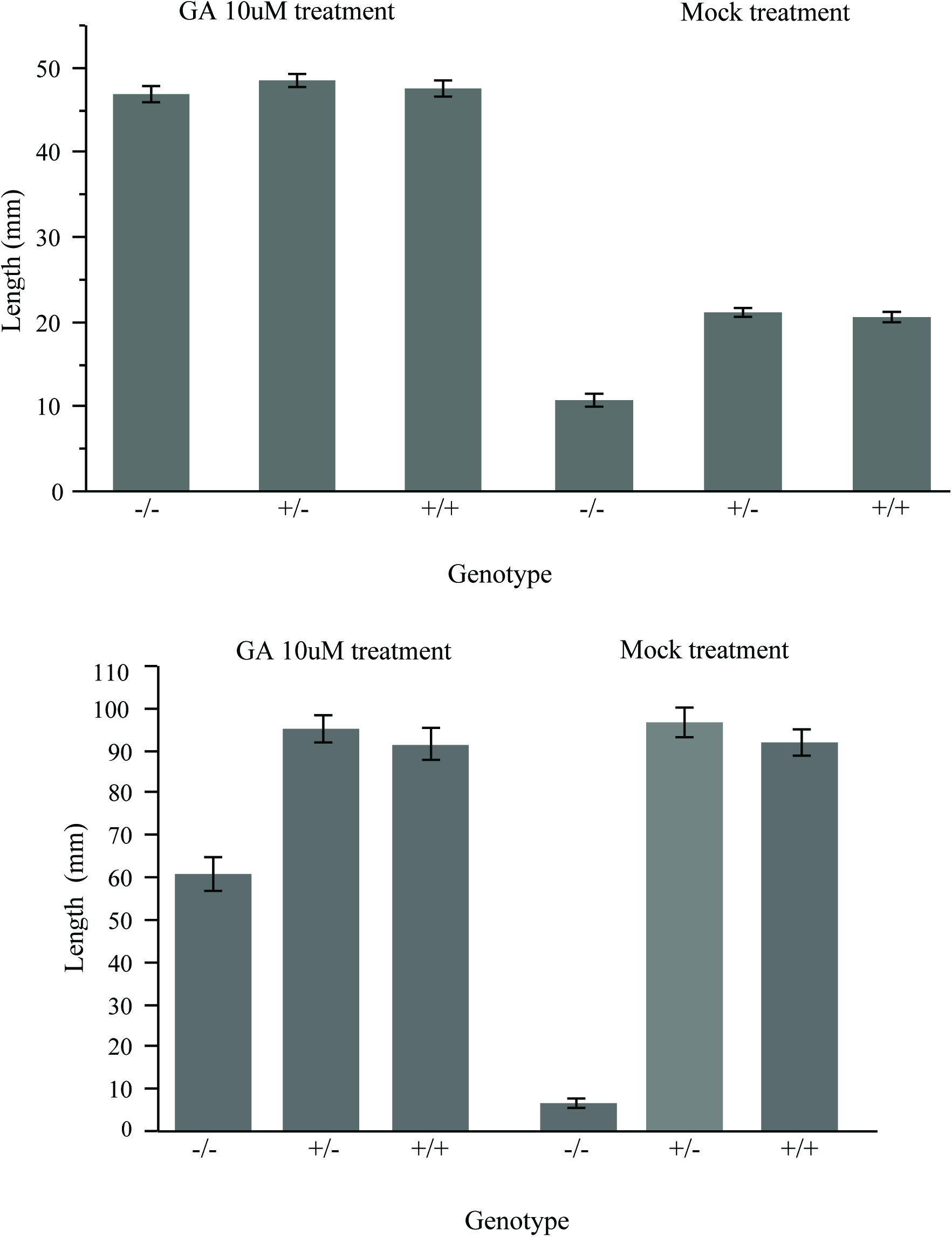
Exogenous gibberellic acid treatment restores growth to *eko-1* seedlings. (a) Photograph of 7 d light-grown seedlings of wild type and *eko-1* mutants treated with 10uM GA (+) or mock treated control (-). (b) Histogram of mean seedling height and standard error for 7 d light-grown seedlings of wild type and *eko-1* mutants treated with 10uM GA (+) or mock treated control (-). (c) Histogram of mean seedling height and standard error for 7 d dark-grown seedlings of wild type and *eko-1* mutants treated with 10uM GA (+) or mock treated controls (-).

### EMS Variation Induced Lines for mutant mapping without segregation variance

We identified the causative *eko-1* polymorphism using recursively selfed, but still segregating (M4), mutagenized material. The SNP calling approach and removal of false positives by comparing our mutant to other lines improved NGS processing for EMS-mutant cloning, increased our confidence in SNP calls, and permitted using EMS-induced variants as molecular markers for mapping (Figures 5 and 6). In addition, by sequencing pools of pedigreed materials we could better distinguish causative polymorphisms from inconsequential background mutations (Figures 1B, 5, 6, and 7). Our success suggested to us that a forward genetics crossing scheme and experimental design that takes advantage of comprehensive information available from whole-genome sequencing was worth exploration.

All sequenced samples were derived from EMS-treated BTx623 plants. Two of these, (10-2 and 12-2) were selected because they had no visible morphological defects. As a result, we have discovered thousands of EMS-induced SNPs that can be used as molecular markers in mapping novel genetic variation without impacting whole-plant phenotype. In addition, the *dhr2-1* mutant has a recessive and screenable blue/white Cu+ reduction phenotype (Krothapalli et al., 2013) that can be used to confirm F1 cross success when the *dhr2-1* mutant is used as the seed parent. The thousands of EMS mutants segregating in that line could then be used as molecular markers in F2 populations derived from confirmed F1 crosses. In order to maximize the utility of these materials, we identifed the SNPs unique to each of the 10-2, 12-2, *dhr2-1* lines. This resulted in 2,288-8,050 SNPs that can be used for mapping (Table 5 and Supplemental Files S1-S3). The two phenotypically unaffected lines in the BTx623 background enable the mapping of mutants of small effect as well as traits strongly affected by the genetic variation between sorghum lines. A cross between these phenotyically normal EVIL twins and any independently-derived mutant provides sufficient molecular markers for mapping without also introducing phentoype-altering segregational variance.

**Table 5.** Summary of EMS Variation Induced Lines for mapping in Sorghum without segregation variance.

| Line | Phenotype | EVIL SNPs |
| --- | --- | --- |
| 10-2 | Unaffected | 8,050 |
| 12-2 | Unaffected | 2,298 |
| *dhr2-1* | Low Cyanide release | 2,288 |

## DISCUSSION

While it would be tempting to conclude from these results that any mutant can be cloned just by sequencing a few lines, there are three essential components to our study that were combined to make it effcient and accurate. First, to limit the mapping procedure to actual polymorphisms it is neccessary to remove all error prone positions in the genome.

Second, we were able to impose logical expectations for SNPs linked to the phenotype that ruled out ~98% of the genome using a pedigree containing a true wild type (no dwarf progeny), a segregating sibling, and a dwarf mutant. If a segregating pedigree is not available, using the EMS-induced SNPs as markers still has many advantages for mapping. The largest advantage is that due to their scarcity, far fewer recombinations will be neccessary to reduce the possible mutation space to a manable list. Finally, the nature of the mutagen used allowed us to narrow down the searchspace to the point where we had one and only one causal locus. The overwhelming majority of the mutations caused by EMS are G to A (with its complement C to T). Furthermore, SNPs affect phenotypic alterations in mutagensis experiments via disruptions in protein-coding sequences as very few single SNPs within non-coding portions of the genome can have large effects on phenotype. Thus, the causal SNP for a mutant phenotype in an EMS-mutagenesis experiment can be identified among G:C to A:T changes that alter protein sequence, as previously discussed (Krothapalli et al., 2013). Other mutagens are more likely to cause insertion/deletion polymorphisms which can affect gene function while acting in non-coding regions (for example by deleting the promoter). Insertion/deletion polymorphisms are also much more difficult to detect using short read sequencing, compounding the problem. It is only with all three factors together that we are able to make our assignment of function.

It is also important to note that getting full coverage of the non-error-prone portion of the genome is an important requirement for this approach, which relies on full coverage of all potential protein coding mutations. False negative SNP calls will lead to the misidentification of SNPs that are identified as being more likely to be the one and only one remaining SNP in the window. In Arabidopsis, simulated experiments estimate a genome coverage of 20-50 times is optimal for NGS mapping, depending on populaiton structure (James et al., 2013). Polyploid genomes, where paralogs may not get separated easily, will also be difficult to analyze using this approach (Tsai et al., 2013).

Fully sequenced lines from the wild-type pedigree (10-2 and 12-2) have great utility for future mapping studies. The lines have no obvious phenotypes and a defined set of real polymorphisms, which make them ideal crossing partners for mapping mutants from mutagenesis types that cannot use the methods described here. In reference to a common theme in literature and popular culture, we have named these lines EVIL twins (EMS Variation Induced Lines) to denote their status as phenotypically identical but genetically different to the wild type. The advantage of crossing to an EVIL twin line is that the only phenotypes that will be segregating in the population are those that are present in the mutant. This could be especially important for cloning of mutants with small effects, in which the phenotypic diversity of the outcross lines can obscure the signal. In addition, radiation mutagenesis induces a wide variety of deletion/insertion polymorphisms along with SNPs, so many of the logical filters we applied here will not work to reduce the false positive calls. Whereas crossing a radiation-derived mutant to a congenic EVIL twin can be used to introduce markers for mapping without causing additional segregation variation. Three of the lines from this study are candidate EVIL twins: 10-2, 12-2 and the *dhr2-1* line. From our analysis, each of these lines has over 2000 unique SNPs that can be used for mapping (Table 3).

A number of called variants visible in Figure 6 are present at frequencies that are not possible for a single sequenced individual. These variant positions depart from the axis of all other SNPs linked to them and appear to defy Mendelian genetics. This pattern was also observed in allele frequency counts from the other genotypes (data not shown). We suspect that these positions represent false positives from error-prone positions that were identified as homozygotes in only one of the lines. When the allele frequencies of these positions are then tallied from the reads data and graphed, we observe an allele frequency that is highly unlikely both due to the proportion and due to the absence of any recombination breakpoints. This is exactly what would be expected if the SNP filtering procedure thresholds an underlying error distribution that is non-randomly distributed across the genome. The subtraction of error prone positions from the five sets of sequencing data presented here are capable of identifying many, but not all, of the false positive positions present in the sorghum genome. Further sequencing experiments by the community will continue to identify positions prone to false positive errors. Indeed, preliminary analysis of a larger set of sequenced lines indicate that additional moderate-frequency shared variants are present in sequence data sets derived from BTx623 and that their discovery was not completely saturated after sequencing 500 mutant lines (Addo-Quaye and Dilkes, unpublished data).

The identified gene responsible for the dwarf allele encodes an enzyme that is the target of Paclobutrazol inhibition of GA biosynthesis (Hedden and Graebe 1985). Mutations of the ortholog of this gene are responsible for the *Gibberellic Acid 3* dwarf mutant of Arabidopsis, the *lh* dwarf mutant of Pea, and the semi-dwarfing allele from Tan-Gibozou rice which contributed to the “green revolution” in Japan (Itoh et al. 2004). Of the maize mutations affected in GA biosynthesis none have yet been recovered for blocks in these steps. The maize genome encodes two of these enzymes resulting from a tandem duplication expressed at dramatically different levels (Supplemental Information). If both are functional, this would explain why this step has not been identified mutationally in maize.

Recently, other studies have found gene mutations in the gibberellic acid and brassinosteroid biosynthesis pathways that impact not only plant height but other agronomically important traits in sorghum. Petti and co-workers (2015) identified a dwarf sorghum mutant, *dwf1-1* that mapped to a frame-shift in a presumed GA20-oxidase. In addition to dwarfism this also caused male sterility and impacted cellulose biosynthesis. This mutation occurs later in the GA-biosynthetic pathway than the *ent*-kaurene oxidase mutation identified in this study. Ordonio et al (2014) identified four loss-of-funciton mutations (ent-copalyl diphosphate synthase (CPS; SbCPS1), ent-kaurene synthase (KS; SbKs1), ent-kaurene oxidase (KO; SbKO1), ent-kaurenoic acid oxidase (KAO; SbKAO1) early in the GA biosynthetic pathway that lead to severe dwarfism and culm bending in sweet sorghum. Finally, Rizal and co-workers (2015) found that a mutation in a cytochrome P450 (CYP90D2) in brassinosteroid bisynthesis lead to decreased vein density in sorghum.

The application of EMS mutagensis and cost effective short read sequencing is driving rapid progress in our understanding of the control of crop plant architecture. Additional improvements in experimental design can further accelerate this and extend the utility of these methods. If we can reliably detect all of the sequence polymorphisms in a given genome, whole genome sequencing allows us to invert the typical process for mutant identification. When all possible causative SNPs are known, only recombinants distinguishing the putatively causative SNPs are needed. This reduces the number of individuals necessary to identify one and only one potentially causative polymorphism in a line. In addition, by using all of the EMS-induced alleles in a mutagenized line, even those in non-coding sequence, we can remove the need to cross a phenotypically selected mutant to a genetically distinct stock. No incidental segregating variance needs to be introduced and, thus, environmentally labile or adaptively consequential mutant phenotypes that would be masked by background variation are now amenable to cloning. Moreover, because there are few mutations of putative effect, and less segregation variance that might drown out the effects of induced alleles with small phenotypic consequences, we can use this approach to quickly and cost-effectively clone mutations that have moderate to small effects.

### Conclusions

Using a novel combination of full genome sequencing, experimental design and bioinformatics algorithms, we have cloned the gene for an EMS-induced dwarf mutant in *Sorghum bicolor*. By comparing the sequence of several lines within the pedigree, removing error prone positions and focusing on polymorphisms that EMS will induce, we were able to narrow the polymorphisms down to one and only one mutation, an induced stop codon in the cytochrome P450 gene Sobic.010G172700. We have demonstrated that this approach can change the current paradigm for mapping mutants. We have also produced three lines that can be used as EVIL twins for mapping in the BTx623 background without introducing additional phenotypic variation.

## MATERIAL AND METHODS

### Plant material

All plant material was grown in the Purdue Horticulture and Landscape Architecture greenhouse in an equal mixture of Turface (Profile Products LLC, Buffalo Grove, IL), potting soil (Conrad Fafard Inc., Agawam, MA), and sand (U.S. Silica, Frederick, MD). For next generation sequencing, whole leaves were collected approximately 1 week after germination for a pool of 10-2 individuals, in which all individuals planted were of normal size and 10-dwarf individuals, in which leaves from all dwarfs germinated across all individuals were collected and pooled. DNA genotyping leaf tissue was collected approximately six weeks after germination from multiple individuals. Leaves from 10-3 were collected from only tall individuals in a segregating population, which included dwarf individuals. Leaves from normal-height 12-2 individuals were collected from a population segregating for a dominant tall variant (data not shown). The *dhr2-1* library was previously described and generated from a single individual (Krothapalli et al., 2013).

### Co-segregation test for the causative dwarf mutation

A dCAPS (Derived Cleaved Amplified Polymorphic Sequences; Neff et. al 2002) were designed for the genomic position 50,477,994 on chromosome 10 using the Washington University in St. Louis dCAPS interface (http://helix.wustl.edu/dcaps/). Primers for PCR amplification were: dCAPs_BstN1-F (5’-TGTGGAAGAGTATGGGAAGGTT-3’) and dCAPs_BstN1-R (5’-CCTCTCCACGTGCAATTCTT-3’). Post PCR-amplification, 10 uL of the PCR product was digested using 5 units of BstN1 (New England Biolabs, Catalog #R0168S), 1X NEB

Buffer 3.1 in a 25 uL total reaction. Digestions were carried out overnight at room temperature and examined using a 2% TAE agarose gel. BstN1 digestions only occurred if PCR amplicon was wild type for the causative SNP, which was 44 bases smaller than the uncut amplicon for individuals with the mutation. Individuals were either wild type (183 base pair band), dwarf (227 base pair band), or heterozygous (both 183 and 227 base pair bands).

### Next-generation DNA sequencing

We generated whole genome sequencing data for the five EMS-treated sorghum individuals using Illumina sequencing technology. The individuals 12-2 and 10-3 were sequenced using an Illumina GAIIx instrument, while the 10-2, 10-d individuals were sequenced using an Illumina Hiseq instrument. Sequenced reads from the 12-2 sample were 80 bases long and single-ends, while the remaining sequenced samples were 100 bases long and paired-ends.

### Sequenced reads mapping

We used BWA version 0.6.2 (Li & Durbin, 2009) to align the next generation sequencing data to the BTx623 sorghum reference genome sequence (version 2.1; Patterson, 2009) which was downloaded from Phytozome (Goodstein et al., 2012). Suffix array co-ordinates for each query sequence were generated using the BWA *aln* command with non-default parameters: “-t 8”. Paired end alignments were generated using the BWA *sampe* command with non-default parameters: “-P ‐r ʺ@RG\tID:SampleID\tSM:SampleName\tPL:Illuminaʺ”. In the case of the single-end reads, sequence alignments were produced using the BWA *samse* command. Sequencing data and mapping statistics were obtained using the SAMtools *flagstat* command with default parameters. Genome coverage statistics were generated from the sequence alignment files using the combination of SAMtools and BEDTools (Quinlan & Hall, 2010). We used the SAMtools *view* command to uncompressed BAM files and sent the results to standard output. We then used the BEDTools *genomeCoverageBed* command to estimate the coverage depth from the uncompressed alignment BAM output. The SAMtools *view* command parameter options were “-u -q 0”, while the BEDTools *genomeCoverageBed* command parameter options were “-ibam stdin ‐g ”.

### SNP calling and quality filtering

SNP detection was performed using the *mpileup* command in the SAMtools package version 0.1.18 (Li et al., 2009). The non-default options invoked were “-B ‐Q 20 ‐P Illumina ‐C50 ‐uf’ and the result was piped to the BCFtools *view* command with non-default parameters: “-vcg”. These settings require bases to have a quality score of 20 or higher, downgrade the consideration of base calls from poorly mapped reads, and disable the base-alignment quality scoring function in SAMtools. Further quality filtering was performed as follows. Removal of SNPs residing in repetitive regions was performed using the *varFilter* command within *perl* script *vcfutils.pl varFilter* with options “-D100” to remove SNPs with coverage depth greater than 100 or less than two. Other default settings for this command also removed SNPs with root mean square quality of less than 10 and SNPs found within three bases of a gap (R. Li et al., 2009). All nucleotide positions for which more than one polymorphism was noted were removed using a custom *awk* script leaving only biallelic positions. Using the SnpSift program (Cingolani et al., 2012a), additional filters were applied. Only homozygous variant positions with a minimum SNP quality of 20 were retained. We required that no reference base reads and at least one observation of the SNP base call from each strand. EMS mutations result in G:C to A:T mutations, whereas false positives could be any change. Thus, we retained only the alleles that correspond to G:C to A:T mutations using SnpSift (See Supplementary Files S1-S5 and Scripts S9-S13 for details).

### Detection of false-positive shared SNPs

Since the three individual plants (10, 12-2 and *dhr2-1*) were obtained from seeds, which were independently treated with EMS, we subtracted all overlapping high quality SNPs detected in the progenies of these individuals. To remove false-positive shared SNPs, we searched for genomic position overlaps in the high quality homozygous G to A, and C to T SNP positions for the five EMS-mutant individuals. For each of the 10-derived individuals (10-2, 10-3 and 10-d) we subtracted the overlaps with the *dhr2-1* and 12-2 individuals. Similarly, we subtracted SNPs in *dhr2-1* with genomic positions overlapping SNP positions in 10-2, 10-3, 10-d and 12-2. The process was repeated for 12-2 by subtracting SNPs with positions overlapping 10-2, 10-3, 10-d and *dhr2-1* SNP positions. We used the BEDTools *intersect* command (version 2.17.0) with options “-wa” to find common SNP variations detected in different samples. Similarly, we used the BEDTools *subtract* command with options “‐A” to detect pedigree- and sample-specific SNPs for each of the samples.

### Allele frequency determination in EMS variation induced lines

For each of the five individuals, we calculated the allele frequencies at all genomic positions corresponding to called variants in each of the other four individuals. To do this, we re-ran SAMtools *mpileup* command and generated a new version of the variant call output files by modifying the BCFtools *view* command non-default parameters to “-Ncg”. The BCFtools *view* command parameter changes results in a new variant call output file that contains genotype calls for both variant and not variant sites, in which the genome reference nucleotide base is known. We used the BEDTools *intersect* command and the coding region information specified in the sorghum genome annotation “.gff3” file, to derive a second set of files containing only the coding region allele information. We then used both the genome-wide and coding region files for each individual and calculated allele frequencies. We subtracted SNPs shared among the members of the 10 pedigree. The BEDTools *subtract* command with options “‐A” was used to detect the subset of high-quality homozygous SNPs that were unique to 10-d and absent in 10-2 and 10-3. These positions were examined in the allele frequency plots to find SNPs that co-segregated with the dwarf phenotype.

### Functional annotation of SNP variation

We used the version 3.1 of the SnpEff program (Cingolani et al., 2012b) to predict the effect of the SNPs on sorghum gene function. We downloaded sorghum version 2.1 whole genome, proteome, coding sequences, and the genome annotation “.gff3” files from Phytozome (version 9.1; www.phytozome.net; Goodstein et al., 2012). These files were used to generate a custom SnpEff database using the SnpEff *build* command with parameter options “build -gff3”. This required editing the “snpEff.config” file to create an entry for sorghum version 2.1. The SnpEff-annotated variant call file was generated using the SnpEff *eff* command with parameter option “-c snpEff.config”. SNPs that altered the amino acid coding capacity of genes were classified as “MODERATE” if they were a non-synonymous change or “HIGH” if they resulted in a nonsense codon, splice loss, or loss of the start codon.

Functional information for the genes affected by these SNPs was obtained as follows. We downloaded the latest proteome sequences and the corresponding annotation files for sorghum BTx623 (version 2.1), maize B73 (version 5b.60) and *Arabidopsis thaliana* (TAIR release 10) from the Phytozome web portal. We created a BLAST proteome databases for each of the three plant species using the *makeblastdb* program with parameter options “-input_type fasta -dbtype prot”. We used the *blastp* program (version 2.2.28+) in the BLAST package (Altschul et al, 1990) to align the soghum proteome to maize and Arabidopsis. The following *blastp* non-default parameter options were selected: “-evalue 1E-05 -num_threads 32 -max_target_seqs 5 -out -outfmt 6 -seg yes”. The sequence annotation for the detected maize and Arabidopsis homologs of the sorghum genes were appended to the SNP call “.vcf” files, using a custom program (appended as supplemental files S6 through S8).

### Dwarf mutant complementation with GA_3_ in the dark

An M2 segregating population of mutant and wild-type siblings was grown in the dark for 7 d at 25°C with 90 % humidity in a Conviron E8 growth chamber (Conviron, Pembina, ND). Seedlings were planted in 72-cell germination flat in a 1:1 mixture of peat germination mix (Conrad Fafard Inc., Agawam, MA) and Turface (Profile Products LLC,

Buffalo Grove, IL). Seedlings were allowed to germinate for 2 d (before emergence) and then were treated with 1.75 L per flat of 0.2% ethanol and 0.005% silwet (“no treatment”) or 10 μM GA_3_ (Gold Biotechnology, St. Louis MO), 0.2% ethanol, and 0.005% silwet (“10 μM GA3 treatment”). Seedlings were harvested and photographed after 7 d total in the dark. Mesocotyl length was measured from the root-to-shoot transition zone to the proximal end of the coleoptile using ImageJ (Abramoff et al, 2012).

### Dwarf mutant complementation with GA_3_ in the light

An M2 segregating population of mutant and wild-type siblings was grown in the light for 14 d under greenhouse conditions: 27°C (day) and 21°C (night) with 16 h supplemental lighting. Seedlings were grown in media as previously stated. Seedlings were allowed to germinate for 2 d then treated with 1 L per flat of 0.2% ethanol and 0.005% silwet (“no treatment”) or 10 μM GA3, 0.2% ethanol, and 0.005% silwet (“10 μM GA3 treatment”) every third day until harvest. Seedlings were harvested and photographed after 14 d total. Plant height was measured from the distal end of the mesocotyl to the first leaf collar using ImageJ (Abramoff et al, 2012).

---

**Supplementary Table 1.** Summary of preliminary and filtered G:C to A:T transitions in the whole genome and protein-coding region SNPs in the error-prone SNPs filtering.

| Line | WG<sup>a</sup><br>gcat% | WG<br>N<sup>b</sup> | CDS<sup>c</sup><br>gcat% | CDS<sup>d</sup><br>N | -<br>BSWG<sup>e</sup><br>gcat% | -<br>BSWG<sup>f</sup><br>N | -<br>BSCDS<sup>g</sup><br>gcat% | -<br>BSCDS<sup>h</sup><br>N |
| --- | --- | --- | --- | --- | --- | --- | --- | --- |
| 10-2 | 49.70 | 19,452 | 45.51 | 3,032 | 77.21 | 10,426 | 69.16 | 1,741 |
| 10-3 | 59.60 | 15,566 | 59.06 | 1,876 | 84.77 | 9,636 | 91.07 | 1,097 |
| 10-d | 50.43 | 18,155 | 44.95 | 2,605 | 78.87 | 9,663 | 71.95 | 1,408 |
| 12-2 | 53.94 | 4,885 | 57.89 | 1,033 | 77.93 | 2,891 | 80.84 | 663 |
| *dhr2-1* | 29.58 | 13,000 | 32.19 | 1,839 | 65.45 | 3,184 | 79.58 | 524 |

<sup>a</sup> WG gcat% refers to Percentage of G:C to A:T SNPs in the whole genome prior to filtering.
<sup>b</sup> WG N refers to Number of G:C to A:T SNPs in the whole genome prior to filtering.
<sup>c</sup> CDS gcat% refers to Percentage of G:C to A:T SNPs in the coding sequences prior to filtering.
<sup>d</sup> CDS N refers to Number of G:C to A:T SNPs in the coding sequences prior to filtering.
<sup>e</sup> -BSWG gcat% refers to Percentage of G:C to A:T SNPs in the whole genome after filtering.
<sup>f</sup> -BSWG N refers to Number of G:C to A:T SNPs in the whole genome after filtering.
<sup>g</sup> -BSCDS gcat% refers to Percentage of G:C to A:T SNPs in the coding sequences after filtering.
<sup>h</sup> -BSCDS N refers to Number of G:C to A:T SNPs in the coding sequences after filtering.

